# Solving scaffolding problem with repeats

**DOI:** 10.1101/330472

**Authors:** Igor Mandric, Alex Zelikovsky

## Abstract

One of the most important steps in genome assembly is scaffolding. Increasing the length of sequencing reads allows assembling short genomes but assembly of long repeat-rich genomes remains one of the most interesting and challenging problems in bioinformatics. There is a high demand in developing computational approaches for repeat aware scaffolding. In this paper, we propose a novel repeat-aware scaffolder BATISCAF based on the optimization formulation for filtering out repeated and short contigs. Our experiments with five benchmarking datasets show that the proposed tool BATISCAF outperforms state-of-the-art tools. BATISCAF is freely available on GitHub: https://github.com/mandricigor/batiscaf.

## 1 INTRODUCTION

Increasing the length of sequencing reads allows assembling short genomes but assembly of long genomes remains one of the most interesting and challenging problems in bioinformatics. Conventionally, the genome assembly process is divided into three stages: contig assembly, scaffolding, and gap filling. Contig assembly refers to the process of building contiguous chunks of DNA from read sequences. The task of scaffolding tools consists of orienting contigs, joining them into chains (called *scaffolds*) and providing distance estimates between neighboring contigs. Significant portions of long genomes are repeated confusing the assembly of the limited length reads which is manifested in numerous contig mis-assemblies and scaffolding errors. Repeats negatively affect scaffolding in two ways: (i) they cause fragmentation of contigs, forcing skipping short contigs and (ii) they produce false connections between non-adjacent contigs, significantly complicating the task of finding the neighboring contigs.

The common strategy to reduce the amount of incorrect joins caused by repeats consists of the following steps (ScaffMatch [10] and BESST [14]):

1. filtering out repeats before the scaffolding process based on their coverage,
2. scaffolding the remaining contigs and
3. inserting some of the filtered contigs in the scaffolding.

Usually the most effort is devoted to step (2), i.e., identification of correct joins between remaining contigs. There are several optimization formulations for step (2), e.g., maximizing the number of “concordant” read pairs (ScaffMatch [10], SILP3 [9], etc) or minimizing the number of “discordant” read pairs (OPERA [4], OPERA-LG [3]). Usually these formulations are NP-complete [4] and solved either heuristically by greedy-like approaches (SSPACE [1], ScaffMatch [10]), or exactly by Integer Linear Programming (ILP) or dynamic programming (OPERA and OPERA-LG).

The first repeat aware scaffolding tool OPERA-LG [3] instead adds as many copies of repeated contigs as necessary. The original OPERA problem formulation of minimizing the number of discordant read pairs is modified to include the repeated contigs in a parsimonious fashion. Thus, OPERA-LG simultaneously scaffolds both unique and repeated contigs. The potential repeats are determined based on the read coverage analysis which may not be accurate enough [3].

It has been recently shown that the state-of-the-art validation framework in [5] does not avoid significant errors in developing scaffolding benchmarks as well as estimating scaffolding quality [8]. Only recently appeared repeat-aware scaffolding evaluation framework (see [8]) allows to compare the quality of repeat-aware as well as standard scaffolding tools which do not specifically handle repeats. Unfortunately, repeat-aware OPERA-LG does not exhibit significant improvement over the original OPERA.

In this paper, we propose a new scaffolding tool *BATISCAF* (BAd conTIg filtering SCAFfolding) which follows the same steps (1-3) but emphasizing filtering out repeats (step (1)) instead of step (2). More precisely, we remove suspected repeats and short contigs which offer multiple alternatives for scaffolding chains. After removal of all confusing contigs, the scaffolding step (2) becomes trivial since no alternatives left. In the final step (3) of inserting removed repeat and short contigs, multiple copies of repeats are added to the appropriate slots between scaffolded contigs.

We have validated BATISCAF on 5 benchmarks: three from the GAGE project [15]: *S. aureus, R. sphaeroides*, and *H. sapiens* (*chr14*) along with two fungi datasets: *M. fijiensis* and *M. graminicola*. Our experimental results show advantage of BATISCAF over existing scaffolding tools.

This paper is organized as follows. The next section contains the description of BATISCAF including the Repeat and Short Contig Filtering Problem formulation with the corresponding ILP formulation and repeat and short contig insertion method. We conclude with the analysis of the experimental results comparing BATISCAF with state-of-the-art scaffolding algorithms.

## 2 METHODS

BATISCAF is a novel repeat aware scaffolding tool. The high-level idea behind it consists of 3 steps:

1. Filtering out potential repeats via ILP
2. Constructing backbone scaffolding for potentially unique contigs
3. Insertion of multiple copies of potential repeats into backbone scaffolds

### 2.1 Repeat and Short Contig Filtering Problem

In this section we describe construction of the scaffolding graph and formulate the repeat and short contig filtering problem. We also show that this problem formulation is NP-hard and can be 3-approximated.

Let *C* be the set of contigs. For each contig *c*_*i*_ ∈*C* we produce two strands – the sequence for the first strand 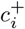 coincides with the contig sequence and the sequence for the second strand 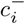 coincides with the reverse complement of *c*_*i*_. We connect the corresponding vertices 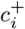 and 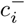 with an *intra-contig* edge 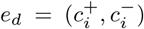 of weight *∞* (a big enough number) (see Figure 1(a)). The set of all intra-contig edges is denoted as *E*_*d*_. Note that each contig *c*_*i*_ ∈*C* has one and only one corresponding intra-contig edge *e*_*d*_ ∈*E*_*d*_.

**Fig. 1.**
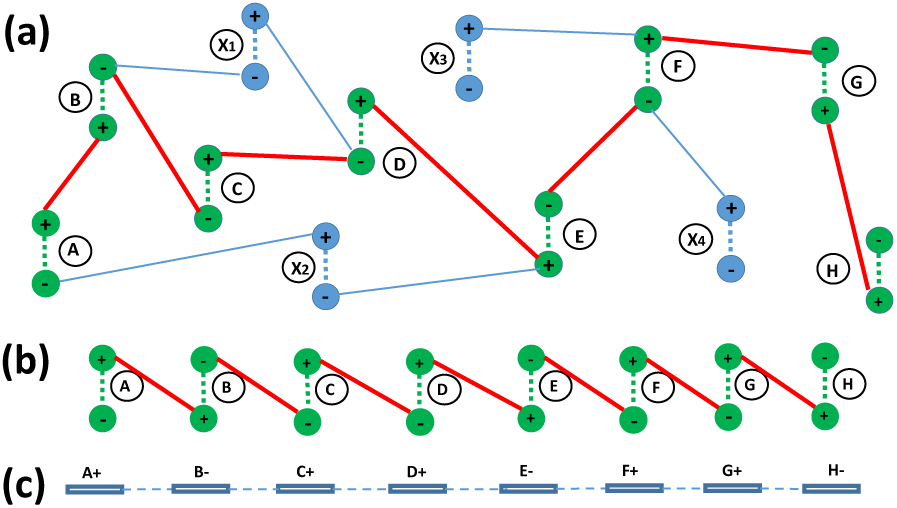
(a) The scaffolding graph *G*. Each contig is represented by two vertices (+ and −) corresponding to forward and backward strands. The intra-contig edges are dashed, the inter-contig edges are solid. (b) The scaffolding graph corresponding to a valid scaffold. The graph is a chain of alternating intra- and inter-contig edges. (c) The chain of contigs corresponding to the scaffolding graph of a scaffold. Each contig is either in the original orientation (+) or inverted (−).

We say that a paired-end read *r* (e.g., Illumina technology) connects strands 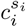 and 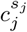 where *s*_*i*_, *s*_*j*_ ∈ {+, −}, *i ≠ j*, of two distinct contigs if the forward read of *r* is aligned to 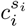 and the reverse read of *r* is aligned to 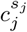. The vertices corresponding to these strands are connected with an inter-contig edge 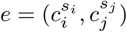 of weight *ω*_*ij*_ if and only if they are connected with *ω*_*ij*_ paired-end reads supporting similar gap estimate (statistically inferred value for the distance in base pairs, gap estimation in BATISCAF follows [13]) between the contigs *c*_*i*_ and *c*_*j*_. Let *E* denote the set of such edges *e*. Finally, the *scaffolding graph G* = (*V* = *C*^+^ ∪ *C*^−^, *E* ∪ *E*_*d*_, *ω*) consist of vertices *V* = *C*^+^ ∪ *C*^−^ corresponding to the contig strands connected with weighted intra- and inter-contig edges.

Any valid scaffold corresponds to a scaffolding graph in which each vertex (strand) is incident to exactly one intra-contig edge and at most one inter-contig edge (see Figure 1(b)). Therefore, if a vertex in *G* is incident to at least two inter-contig edges, then either one or both of them should be disregarded. Assuming no contig mis-assemblies such confusing edges are caused by either repeat or short contigs (see Figure 2). If there is no such confusion, the scaffolding graph *G* is a set of valid scaffolds (with potentially missing links). So in order to avoid confusion and keep only reliable contigs we need to delete suspected repeat and short contigs. Usually, repeat contigs are also short (which is defined by the repeat monomer length of 150-400 bp [11]). Thus, the problem of repeat and short contig removal can be formulated as the following

**Fig. 2.**
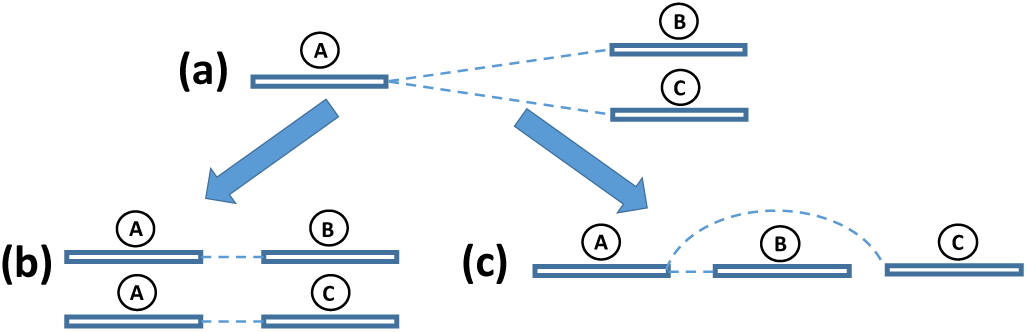
(a) A confusion triple: the same strand of contig *A* is connected with two strands of contigs *B* and *C*; The two possible scenarios causing the confusion: (b) The contig A is a repeat and another copy of contig *A* is connected with *C*; (c) The connection from *A* to *C* jumps over the short contig *B*.

#### Repeat and Short Contig Filtering (RSCF) Problem

Given a scaffolding graph *G* = (*V, E* ∪ *E*_*d*_, *ω*) find minimum total length subset of contigs *X* ⊆ *V* such that in subgraph *G′* induced by *V \ X*, any vertex *v* is incident to at most one inter-contig edge.

The solution of the RSCF problem represents a set of contigs whose removal from *G* leaves a set of simple alternating paths and/or cycles consisting of intra-contig edges interspersed with inter-contig ones. The paths can be easily transformed into scaffolds. If the are no cyclic chromosomes, we need to transform them into paths. Therefore, from each cycle we remove the least weight edge. After the least weight edge is removed from each cycle the resulting alternating paths can be easily transformed into a set of scaffolds using a procedure similar to [10] (see Figure 1(c)). Such scaffolds are highly reliable since there are no confusion during their extraction. Clearly, solving the RSCF problem does not guarantee to exhaustively remove all the repeated contigs from *C*. Indeed, a contig with a high degree of its strands which is not necessarily a repeated one in the scaffolding graph would be a candidate for removal. The objective of RSCF problem ensures that the minimal length contigs are removed.

The RSCF problem can be viewed as a set cover problem in which elements correspond to confusion triples of contigs, i.e., contig triples with two *E*-edges connecting a single strand with two different strands of other contigs (see Figure 2) and sets correspond to contigs – each contig *c* covers all confusion triples containing *c*. Therefore, this is a set cover problem where each element belongs to at most three sets. Although such a problem is NP-complete, it can be 3-approximated with a primal-dual algorithm [16].

The RSCF problem can be solved efficiently if the scaffolding graph does not contain cycles. Therefore, instead of solving this problem exactly or approximately, we also propose a fast heuristic consisting of finding the maximum spanning tree *T* (*G*) of the scaffolding graph *G* (note that edges connecting the strands of the same contig will belong to *T*) and then finding the exact solution for *T* (BATISCAF-MST).

### 2.2 Integer Linear Program Formulation for the RSCF Problem

Since the RSCF problem is NP-hard, we propose to find the exact optimal solution using an Integer Linear Programming (ILP) approach.

Let *G* = (*V, E* ∪ *E*_*d*_, *ω*) be the scaffolding graph, where *V* is the set of contig strands.

Let the length (in bp) of contig *u* be denoted as *l*_*u*_. We formulate the following ILP:

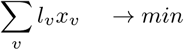

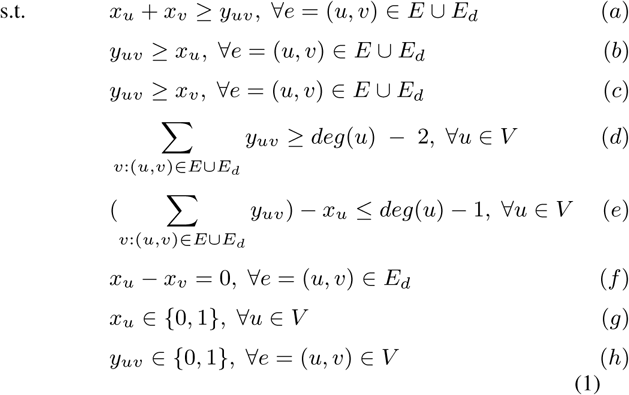

Binary variables *x*_*u*_ ∈ {0, 1} have the following meaning:

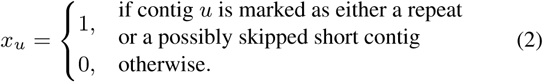

#### Definition

Let *u* be a contig belonging to the scaffolding graph *G*. If by solving the ILP (1) we obtain *x*_*u*_ = 1 we call such contig *untrusted* (otherwise, *trusted*). The set of all untrusted contigs is denoted as *U.*

Binary variables *y*_*uv*_ ∈ {0, 1} have the following meaning:

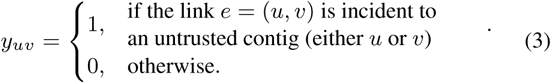

#### Definition

Let *e* = (*u, v*) be an edge in the scaffolding graph *G*. We call the edge *e untrusted* if it is incident to at least one untrusted contig.

The meaning of all the constraints is the following:

- Conditions (a), (b), (c) from the ILP (1) ensure that edges incident to an untrusted contig are also marked as untrusted.
- Condition (d) means that the degree of a trusted contig *u* is at most 2 in the resulting scaffolds, i.e. no more than two edges which are not marked as untrusted are incident to it.
- Condition (e) means that if all the edges incident to a node *u* are marked as untrusted than *u* is untrusted as well.
- Condition (f) guarantees that two strands of any contig are both are either marked as trusted or untrusted.

Finally, the objective of the ILP (1) is to minimize the total length of the untrusted contigs while requesting that all the trusted contigs be connected into chains or cycles (i.e. their degree in the graph *G* \ *U* is either 0, 1, or 2).

The solution of ILP (1) represents a set of untrusted contigs *U.* After we remove them from the scaffolding graph we obtain a graph *T* = *G \ U* with all its connected components being either paths or cycles. We remove the least weight edge from each cycle. As a result, we obtain a set of alternating paths which can be translated into scaffolds following a procedure similar to [10]. In each scaffold *S* relative ordering and orientation of each contig is established.

### 2.3 Second Stage. Most Likely Repeat and Short Contig Insertion

The second stage of our algorithm which is constructing scaffolding corresponding to contigs left after removing confusing contigs (see Figure 1(c)) is trivial. The third stage of our approach is to insert the removed contigs from the set *U* back into the scaffolds. Potential repeats identified in the first stage are inserted into the scaffolds as many times as it can be inferred from the scaffolding graph structure. For each scaffold *s* ∈ *S* we create a *surrounding* graph *G*_*S*_ = (*S* ∪ *N, E*) which is a subgraph of the scaffolding graph *G* on the set of nodes representing contig strands in *S* and all their neighboring nodes *N,* i.e. ∀*n* ∈ *N*, ∃*u* ∈ *S*, such that *e* = (*n, u*) ∈ *E*(*G*) (see Figure 3a). In *G*_*S*_, the orientation of each contig in *S* is known and the relative order between contigs in *S* is established. The same information is to be determined for the contigs in *N.* Surrounding graphs *G*_*s*_*i* corresponding to different scaffolds *s*_*i*_ ∈ *S* may share neighboring nodes, i.e. 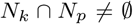 for some scaffolds *s*_*k*_ and *s*_*p*_. This fact determines the copy number of each repeated contig.

**Fig. 3.**
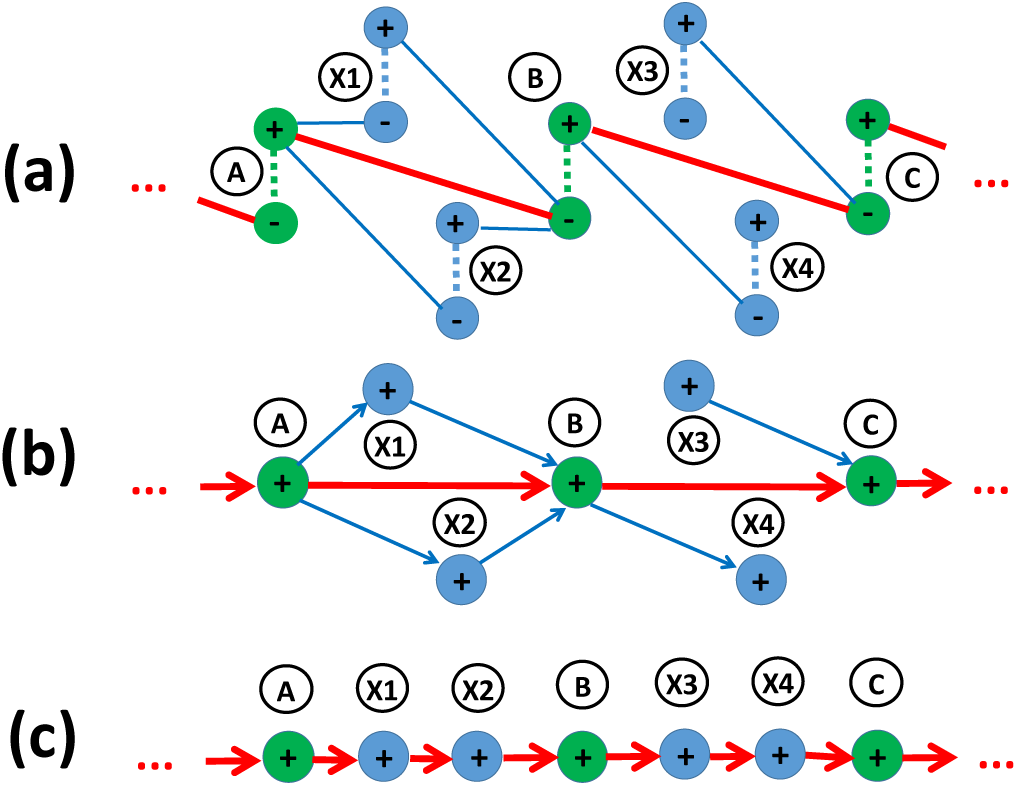
Insertion procedure. Contigs belonging to a backbone scaffold *S* have green color; contigs which are candidates for insertion have blue color.a) A fragment of surrounding graph 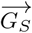 with the chain of trusted contigs *A, B, C* and neighboring contigs *X*_1_– *X*_4_. b) The directed surrounding graph 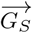 corresponding to *G*_*S*_. c) The scaffold *S* augmented with contigs *X*_1_, *X*_2_, *X*_3_, and *X*_4_.

Next, we build a directed surrounding graph 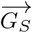 where nodes represent oriented contigs and arcs encode the relative ordering of neighboring contigs. The orientation of each contig in *N* as well as the relative ordering between any contig *n* ∈ *N* and its neighbors from *S* in *G*_*S*_ can be determined in the following way. Let *e* be an edge in the surrounding graph *G*_*S*_ between a strand of contig *n*∈ *N* and a strand of a trusted contig *u* ∈ *S*. If orientation of *u* is determined to be “+” and *e* is incident to the negative strand of *u* then if *e* is also incident to the positive strand of *n* the orientation of *n* is assigned to be “+” (otherwise “–”). A new arc from *n* to *u* is added to 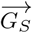 In the same manner, the direction of each arc and the orientation of each contig is determined. For example, in the graph *G*_*S*_ depicted in Figure 3a, the positive strand of the contig *X*_1_ ∈ *N* is connected to the negative strand of contig *B* ∈ *S* (which has orientation “+”). Therefore, in the graph 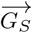 depicted in Figure 3b, contig *X*_1_ is assigned orientation “+” and there is an outgoing arc connecting it with *B*.

The directed graph 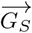 may not be acyclic because some of the newly introduced contigs (from the set *N*) into the scaffold are repeats. We have to identify the set of repeated contigs *R* and for each contig *r* ∈ *R* we replace it with several copies of itself. The resulting graph is acyclic, i.e. it represents a partial order. A minimal set of repeated contigs in 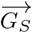 can be determined by solving the Minimum Feedback Vertex Set problem or any of its weighted versions (for example, using contig lengths as weights). It is known that this problem is APX-hard on directed graphs [2], i.e. it does not admit any polynomial time approximation schemes (PTAS). Therefore, we apply a simple greedy heuristic to determine the feedback vertex set. Namely, we randomly pick a cycle 𝒞 in 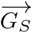, find the smallest length contig *c* ∈ 𝒞 and remove *c* from the graph. We assign a copy of *c* to each of its neighbors in 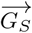 We continue this procedure until the graph is acyclic.

We refer to the transitive reduction of the directed acyclic graph (DAG) 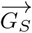 as *spine*. The spine consists of all the nodes in the scaffold *S* and some or all the *N* nodes.

We define a *slot* 𝒮= (*u, v*) as a set of nodes between a pair of articulation nodes *u* and *v* in the spine of 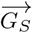 which does not contain any other articulation point. For a slot 𝒮 there can be only two cases (or a combination of them):

1.It is composed of a set of directed paths from *u* to *v* (e.g., in the Figure 3b, the slot (*A, B*) comprises two paths: *A → X*_1_ *→ B* and *A → X*_2_ *→ B*);

2. It contains only “dangling” nodes attached either to *u* or to *v* (e.g., in the Figure 3b, the slot (*B, C*) contains a dangling node *X*_4_ attached to *B* and another dangling node *X*_3_ attached to *C*);

In the first case, for all the contigs belonging 𝒮 to we identify their relative ordering by sorting them according to the distance from either *u* or *v*. In the second case, a “dangling” node 𝒟 may not necessarily belong to. For example, in Figure 3b, contig *X*_3_ may belong to either slot (*B, C*) or (*A, B*) depending on the distance estimates between *X*_3_ and *C* and between *B* and *C*. Without loss of generality, let’s consider 𝒟 to be connected with a contig *S*_*i*_ ∈ *S* by an outgoing arc (e.g. contig *X*_3_ has an outgoing arc to *C*). Contig 𝒟 may be inserted into one of the slots (*S*_*i*–1_, *S*_*i*_), (*S*_*i*–2_, *S*_*i*–1_), etc. For all such slots 𝒮 _*k*_ = (*S*_*i*–*k*–1_, *S*_*i*–*k*_), we estimate the probability 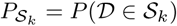 defined as

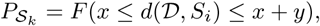

Where

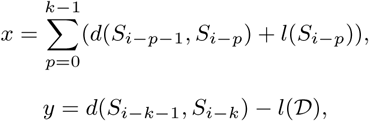

*l*(*z*) is the length of contig *z, d*(*z*_1_, *z*_2_) is distance estimate between two contigs *z*_1_ and *z*_2_, *F* is the normal distribution *N* (*µ, σ*^2^) with *µ, σ*^2^ being the mean and standard deviation of the library insert size.

The dangling contig *d* is assigned to the slot 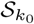, where 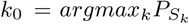. After all the contigs in *N* are assigned to the corresponding slots, we get the set of scaffolds *S′* augmented with repeated and short contigs (see Figure 3c).

## 3 RESULTS

In this section we are aimed at comparing our tool with most of the standard scaffolding tools and in particular to contrast it with OPERA-LG. In our comparison we are concerned with building such a dataset in which every contig has an integer copy number, i.e. there are no partial repeats. In silico simulated datasets where contigs are produced as random DNA sequences with integer copy numbers is an easy task. However, such a dataset would not resemble a real one. For such “perfect” contigs any scaffolding tool performs well [5]. To create a dataset which would be as “close” as possible to a real one and in the same time each contig would be repeated an integer number of times is hard due to mosaic structure of sub-repeats [12] in genomes. We developed a procedure for generating datasets where each repeat is represented by a separate contig.

### 3.1 Building datasets with repeats

Following [5], we align the contigs produced by the Velvet assembler [17] to the reference genome using nucmer [6]. But unlike [5], we consider all the nucmer alignment hits with *α*-identity (we used *α* = 97%) including partial alignments of at least *λ* base pairs long (we used *λ* = 200). Then we split the alignments into segments defined by alignment endpoints (see Figure 4). The segments of length less than *λ* bp are dropped. The remaining segments make artificial contigs. Then multiple artificial contigs are collapsed into a single artificial contig if the corresponding genome sequences are *α*- identical. The produced artificial contigs are either repeats or unique sequences at least *λ* bp long. Note that no artificial contig contain *λ*-long repeats.

**Fig. 4.**
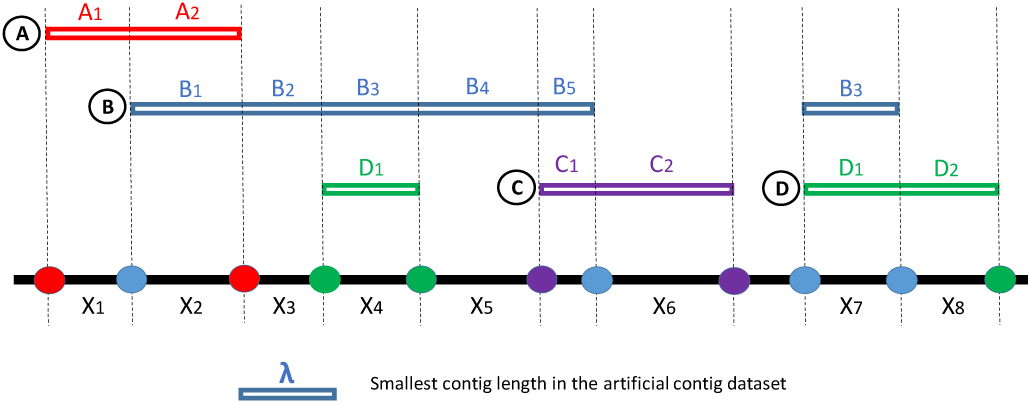
Original contigs *A, B, C* are aligned to the reference with nucmer. Contig *A* significantly overlaps with contig *B*, contig *B* overlaps with contig *C*. Contig *B* contains a repeated region *B*_3_ which also aligns to a different place on the reference. Overlap length of *A* and *B* (which is *A*_2_*≡B*_1_ *≡X*_2_) is greater than the minimum contig length parameter *λ*, therefore we retain *X*_2_. Overlap of *B* and *C* (which is *B*_5_*≡C*_1_) is not included in the artificial contig dataset because its length (length of sequence *B*_5_*≡C*_1_ is smaller than the threshold *λ*. Contig *D* has a partial alignment to *X*_4_ as well as contig *B* has a partial alignment to *X*_7_. The two segments *X*_4_ and *X*_7_ are collapsed into a single artificial contig *X*_4_. Finally, the artificial contig dataset consists of *X*_1_, *X*_2_, *X*_3_, *X*_4_, *X*_5_, *X*_6_, *X*_8_.

**Fig. 5.**
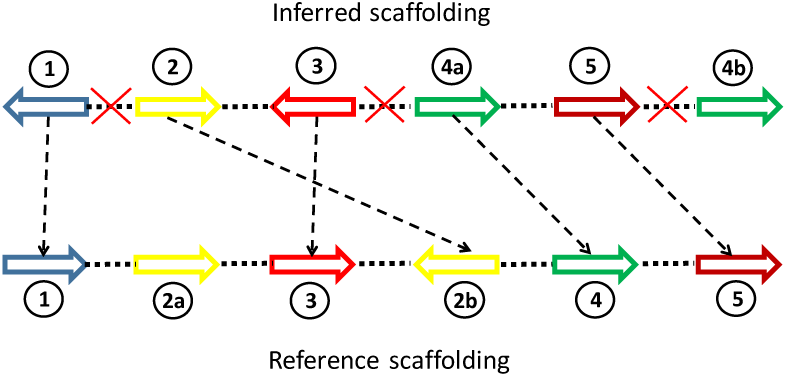
Matching of contigs in the scaffolding *S* and in the reference *R*. Only the links (2,3) and (4a,5) are mapped correctly. Contig 2 has two copies 2a and 2b in the reference scaffolding, contig 4 is inferred to have two copies 4a and 4b. Assigning contig 2 to either of the copy 2a or 2b, as well as assigning either 4a or 4b to the reference contig 4 affects the number of correctly inferred contigs links. Indeed, assigning contig 2 to the reference contig 2a and contig 4b to 4 will mistakenly undercount the number of correct links. The optimal assignment (2 to 2b, 4a to 4) allows to detect two correctly linked contig pairs [8].

We built two sets of artificial contigs based on Velvet assembly from the GAGE project: *S. aureus, R. sphaeroides*, and *H. sapiens* (*chr14*). Also, we included two novel artificial (generated with the aid of iWGS pipeline [18]) fungi datasets: *M. fijiensis* and *M. graminicola*.

The following Illumina paired-end read datasets were used: *S. aureus* - read length 37, insert size 3600; *R. sphaeroides* - read length 101, insert size 3700; *M. fijiensis* - read length 100, insert size 1800; *H. sapiens* (*chr14*) - read length 101, insert size 2700; *M. graminicola* - read length 100, insert size 1800. In the Table 1 the basic characteristics of the contig datasets are presented.

**Table 1.** The basic characteristics of the simulated contig datasets: “avg len”- average contig length, “# unique” - the number of unique contigs (no copies counted), “# total” - the total number of contigs including copies, “# repeats” - the number of repeated contigs, “# max CN” - the maximum copy number, i.e. the number of times the most abundant contig is encountered in the dataset.

|  | avg len | # unique | # total | # repeats | # max CN |
| --- | --- | --- | --- | --- | --- |
| *S. aureus* | 13.9 K | 203 | 244 | 22 | 5 |
| *R. sphaeroides* | 7.2 K | 612 | 652 | 30 | 5 |
| *H. sapiens* (*chr14*) | 1.7 K | 44350 | 45035 | 613 | 5 |
| *M. graminicola* | 4.7 K | 6875 | 7261 | 280 | 9 |
| *M. fijiensis* | 2.0 K | 17781 | 19357 | 995 | 28 |

### 3.2 Evaluation framework

The standard scaffolding evaluation framework described in [5] is not suitable for analyzing performance of repeat-aware scaffolding tools as it takes into account only one possible placement for each contig in scaffolds. For a repeat-aware evaluation, we used a new framework described in [8] which is available at https://github.com/mandricigor/repeat-aware.

In this framework, the contigs from an “inferred” scaffolding (output of a scaffolding tool) are optimally matched with the contigs from the reference scaffolding for which relative order, orientation and distance between neighboring contigs are known. The optimality criterion is based on the number of correct links matched between the two set of scaffolds. This framework allows comparing both standard and repeat-aware scaffolding tools.

### 3.3 Performance metrics

We used the following evaluation metrics in our comparison:

1. number of correct contig links as output by the framework described in Section 3.2;
2. sensitivity (or recall) and PPV (positive predictive value) - two scaffolding quality metrics introduced in [10] and used in [7]. They are defined as 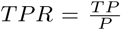, 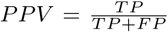, where *TP* is the number of correct contig joins in the output of the scaffolder (true positives), *FP* be the number of erroneous joins (false positives), and *P* is the number of potential contigs that can be joined in scaffold (equal to the number of contigs minus the number of reference sequences). We also report F-score equal to the harmonic mean of *TPR* and *PPV*;
3. Corrected N50 which is the length of contigs in the smallest corrected scaffold necessary to cover 50% of all contigs [15].

### 3.4 Validation results

We compared our tool with other five state-of-the-art stand-alone scaffolders: OPERA-LG, SSPACE, BESST, ScaffMatch, and BOSS. On each of the five datasets BATISCAF outperforms all other tools (Table 3). Notably, a large gap between BATISCAF and the remaining tools is observed on the GAGE datasets *S. aureus, R. sphaeroides*, and *H. sapiens* (*chr14*) in terms of both F-score and corrected N50 metrics.

**Table 2.**
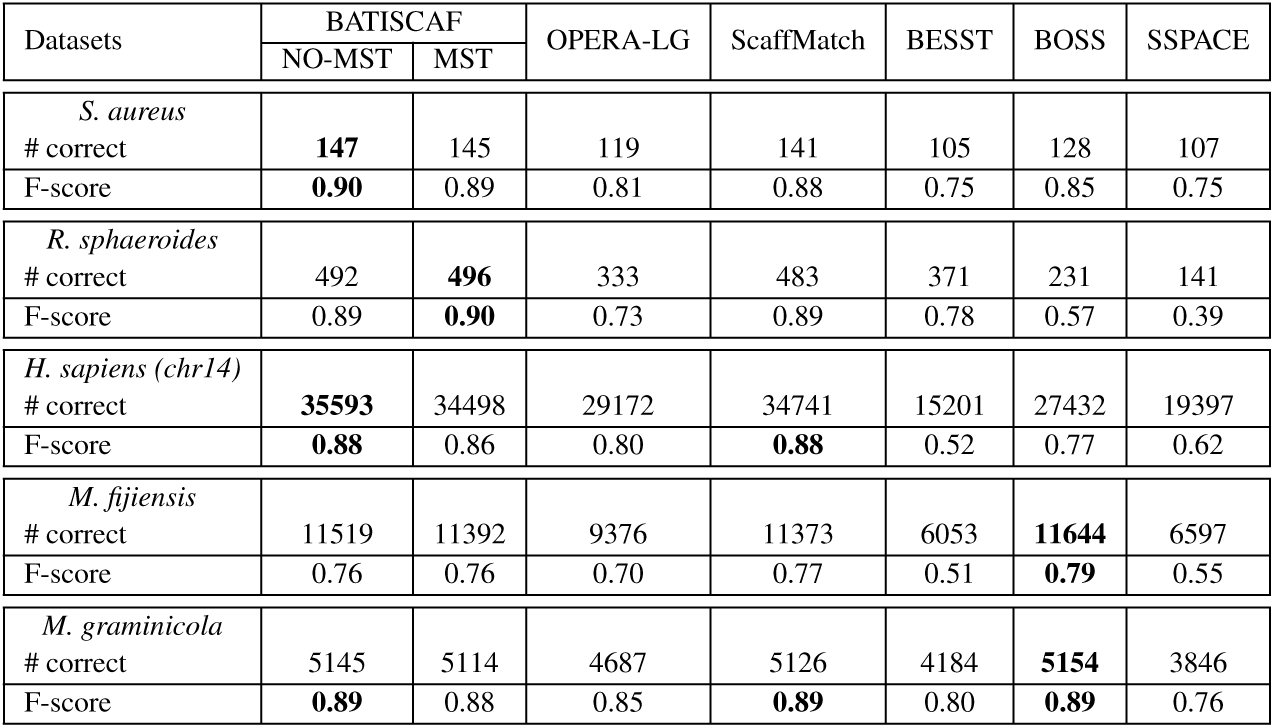
The evaluation results using the standard repeat unaware framework [5].

| Datasets | BATISCAF |  | OPERA-LG | ScaffMatch | BESST | BOSS | SSPACE |
| --- | --- | --- | --- | --- | --- | --- | --- |
|  | NO-MST | MST |  |  |  |  |  |
| *S. aureus* |  |  |  |  |  |  |  |
| # correct | **147** | 145 | 119 | 141 | 105 | 128 | 107 |
| F-score | **0.90** | 0.89 | 0.81 | 0.88 | 0.75 | 0.85 | 0.75 |
| *R. sphaeroides* |  |  |  |  |  |  |  |
| # correct | 492 | **496** | 333 | 483 | 371 | 231 | 141 |
| F-score | 0.89 | **0.90** | 0.73 | 0.89 | 0.78 | 0.57 | 0.39 |
| *H. sapiens (chr14)* |  |  |  |  |  |  |  |
| # correct | **35593** | 34498 | 29172 | 34741 | 15201 | 27432 | 19397 |
| F-score | **0.88** | 0.86 | 0.80 | **0.88** | 0.52 | 0.77 | 0.62 |
| *M. fijiensis* |  |  |  |  |  |  |  |
| # correct | 11519 | 11392 | 9376 | 11373 | 6053 | **11644** | 6597 |
| F-score | 0.76 | 0.76 | 0.70 | 0.77 | 0.51 | **0.79** | 0.55 |
| *M. graminicola* |  |  |  |  |  |  |  |
| # correct | 5145 | 5114 | 4687 | 5126 | 4184 | **5154** | 3846 |
| F-score | **0.89** | 0.88 | 0.85 | **0.89** | 0.80 | **0.89** | 0.76 |

**Table 3.** The evaluation metrics on the five datasets (*α* = 0.97, *λ* = 200) as obtained from solving the ILP (1). The bold font marks the best results.

| Datasets | BATISCAF |  | OPERA-LG | ScaffMatch | BESST | BOSS | SSPACE |
| --- | --- | --- | --- | --- | --- | --- | --- |
|  | NO-MST | MST |  |  |  |  |  |
| *S. aureus* |  |  |  |  |  |  |  |
| # correct | **172** | 169 | 109 | 129 | 87 | 94 | 105 |
| Sensitivity | **0.71** | 0.70 | 0.45 | 0.54 | 0.36 | 0.39 | 0.44 |
| PPV | **0.85** | 0.84 | 0.69 | 0.70 | 0.66 | 0.62 | 0.76 |
| F-score | **0.77** | 0.76 | 0.54 | 0.61 | 0.47 | 0.48 | 0.56 |
| Scaffold corrected N50 (K bp) | **245.3** | 228.4 | 113.3 | 159.8 | 112.2 | 112.2 | 159.4 |
| *R. sphaeroides* |  |  |  |  |  |  |  |
| # correct | 500 | **501** | 333 | 405 | 358 | 196 | 114 |
| Sensitivity | 0.77 | **0.78** | 0.52 | 0.63 | 0.56 | 0.30 | 0.18 |
| PPV | 0.87 | 0.86 | 0.84 | 0.91 | **0.94** | 0.79 | 0.76 |
| F-score | **0.82** | **0.82** | 0.64 | 0.69 | 0.70 | 0.43 | 0.29 |
| Scaffold corrected N50 (K bp) | **679.2** | 361.5 | 85.7 | 173.1 | 209.8 | 28.5 | 21.2 |
| *H. sapiens (chr14)* |  |  |  |  |  |  |  |
| # correct | **30916** | 30092 | 23168 | 20780 | 12376 | 5280 | 12096 |
| Sensitivity | **0.69** | 0.67 | 0.51 | 0.46 | 0.27 | 0.12 | 0.27 |
| PPV | **0.79** | 0.76 | 0.77 | 0.57 | **0.79** | 0.50 | 0.61 |
| F-score | **0.74** | 0.71 | 0.61 | 0.51 | 0.40 | 0.19 | 0.37 |
| Scaffold corrected N50 (K bp) | **33.4** | 25.1 | 17.4 | 19.4 | 11.0 | 4.3 | 9.9 |
| *M. fijiensis* |  |  |  |  |  |  |  |
| # correct | 10397 | **11019** | 7913 | 10095 | 5442 | 6093 | 5099 |
| Sensitivity | 0.54 | **0.57** | 0.41 | 0.52 | 0.28 | 0.32 | 0.26 |
| PPV | 0.84 | 0.85 | 0.82 | 0.82 | **0.89** | 0.78 | 0.80 |
| F-score | 0.66 | **0.68** | 0.55 | 0.64 | 0.43 | 0.45 | 0.39 |
| Scaffold corrected N50 (K bp) | **26.9** | 25.9 | 16.9 | 24.5 | 15.6 | 11.3 | 11.0 |
| *M. graminicola* |  |  |  |  |  |  |  |
| # correct | 5330 | **5455** | 4490 | 5237 | 4184 | 4354 | 3522 |
| Sensitivity | 0.74 | **0.75** | 0.62 | 0.72 | 0.58 | 0.60 | 0.49 |
| PPV | 0.93 | 0.93 | 0.89 | 0.93 | **0.94** | 0.89 | 0.89 |
| F-score | 0.82 | **0.83** | 0.73 | 0.81 | 0.72 | 0.72 | 0.63 |
| Scaffold corrected N50 (K bp) | **101.9** | 97.5 | 64.8 | 89.2 | 75.7 | 56.2 | 38.2 |

Indeed, on *S. aureus* it identified 33 %, on *R. sphaeroides* 23%, and on *H. sapiens* (*chr14*) 34 % more correct contig links than the next top competitor (ScaffMatch on the first two datasets and OPERA-LG on the third one) (see Table 3).

BATISCAF scaffolds for the fungi datasets are also of a better quality, although it did not aggressively join contigs as on the GAGE datasets. However, a small improvement over ScaffMatch in terms of F-score is compensated by more contiguous scaffolds as it is suggested by the corrected N50 results (see Table 3).

It is worth mentioning that BESST is the top “cautious” scaffolding tool (it has top PPV on 4 out of 5 benchmarks).

The runtime of BATISCAF on the largest *H. sapiens* (*chr14*) dataset containing 44350 distinct contigs is reasonable (≈ 80 minutes) and comparable with runtime of other tools and the wall clock time spent on solving the ILP (1) is 16 seconds. We used CPLEX (version 12.7) for solving the ILP (1). All the experiments were run on 2.5GHz 16-core AMD Opteron 6380 processors with 256Gb RAM running under Ubuntu 16.04 LTS.

Further, we also used the well-established standard evaluation framework [5] to confirm the advantage of BATISCAF over the competitor tools. As we mentioned previously, it does not take into account repeats and chooses only the “best” placement of each contig in the reference ground truth scaffolding. We generated artificial contigs for the same five datasets using the scripts from [5] (https://github.com/martinghunt/ Scaffolder-evaluation). Note, that these contigs are different from the ones we used in the repeat aware evaluation. The results in Table 2 suggest that BATISCAF is a top performer even in repeat unaware settings.

## 4 CONCLUSIONS

We presented a novel highly performing repeat-aware scaffolding tool BATISCAF (and its faster version BATISCAF-MST). Our tool solves the scaffolding problem in an innovative way. Instead of tackling it directly, BATISCAF first solves the problem of minimal length repeat and short contig removal after which the problem of scaffolding becomes trivial. The remaining contigs are connected into very reliable scaffolds which are then augmented with the previously removed contigs. The procedure for insertion of short and repeated contigs into the scaffolds detects the necessary number of times each repeated contig to be inserted. For each contig copy it finds the most likely slot for insertion.

We validated BATISCAF on 5 benchmarking datasets and compared it to other state-of-the-art scaffolders. Our experiments showed that BATISCAF is the top performer in terms of F-score and corrected N50 on all the datasets.

The future work on BATISCAF will be focused on providing the users the possibility to scaffold contigs using multiple libraries (including second and third generation sequencing).

